# Self-generating autocatalytic networks: structural results, algorithms, and their relevance to evolutionary processes

**DOI:** 10.1101/2023.09.01.556005

**Authors:** Daniel Huson, Joana C. Xavier, Mike Steel

## Abstract

The concept of an autocatalytic network of reactions that can form and persist, starting from just an available food source, has been formalised by the notion of a Reflexively-Autocatalytic and Food generated (RAF) set. The theory and algorithmic results concerning RAFs have been applied to a range of settings, from metabolic questions arising at the origin of life, to ecological networks, and cognitive models in cultural evolution. In this paper, we present new structural and algorithmic results concerning RAF sets, by studying more complex modes of catalysis that allow certain reactions to require multiple catalysts (or to not require catalysis at all), and discuss the differing ways catalysis has been viewed in the literature. We then focus on the structure and analysis of minimal RAFs, and derive structural results and polynomial-time algorithms, with applications to metabolic network data described briefly.

## 1 Introduction

A central property of the chemistry of living systems is that they combine two basic features: (i) the ability to survive on an ambient food source and (ii) each biochemical reaction in the system requires only reactants and a catalyst that are provided by other reactions in the system (or are present in the food set). The notion of a self-sustaining ‘collectively autocatalytic set’ captures these basic features, and their study was pioneered by Stuart Kauffman [24, 25]. By investigating a simple binary polymer model, Kauffman showed that collectively autocatalytic sets invariably emerge once the network of polymers becomes sufficiently large.

The notion of a collectively autocatalytic set was subsequently formalized more precisely as a ‘reflexively autocatalytic and food-generated’ (RAF) set (defined shortly). RAF sets (RAFs) are related to, but somewhat different from Robert Rosen’s (M, R) systems (a partial connection between the two was described in [22]). RAF theory can also be investigated within the framework of Chemical Organisation Theory (COT) [4]; for example, certain types of RAFs correspond to chemical organisations, as described in [19] (see also [40] Section 4).

RAF algorithms have also been used in the analysis of simple autocatalytic networks of polymers in laboratory studies, either from RNA molecules RNA molecules [16] or from peptides [15], and have been discussed further in modelling the origin of life (see e.g., [6, 44]). More recently, RAFs have also played a pivotal role in modeling self-reproduction and self-organisation before the emergence of a genetic code in the polymer world. When modelling this early stage of chemical evolution [46, 47], RAFs provide a framework for studying the organisation of small molecules in complex chemical networks that would then lead to polymers as RNA and protein. A general framework to model complex catalysis is required here: multiple small-molecules can be involved in catalysis of the same reaction, whereas some reactions can occur spontaneously. There are many examples of this in biochemistry; for example, the enzyme L-threonine dehydrogenase, important in amino acid metabolism requires an organic cofactor, namely NAD, and a metal [23]. Moreover, in the data set analysed in [46] 1052 of the 5994 biochemical reactions involved catalysts combinations of two or more molecules to be present.

Two important features of the RAF approach are the degree of generality RAFs allow, and the fact that very large systems can be analysed precisely by fast algorithms. The generality of RAF theory means that a ‘reaction’ need not refer specifically to a chemical reaction, but to any process in which certain items are combined and transformed into new items, and where similar items facilitate (or catalyse) the process without being used up in the process. This has led to the application of RAF theory to processes beyond biochemistry, such as cognitive modelling in cultural evolution [7, 8], ecology [9, 10], and economics [10]. This generality is not unique to RAFs; for example, Chemical Organisation Theory has been applied to diverse settings including sociology [5], ecology [11], cybernetics [13] and modelling of worldviews [45]. Petri Nets have also been also been applied to biochemical modelling, including self-reproduction [37] and to other non-biochemical settings (e.g. [27]). In addition, a number of other recent structural approaches to autocatalytic networks have been applied in the origin-of-life setting [2, 35].

In this paper, we describe further extensions and applications of RAF theory. We describe an extension of the RAF approach that provides a unified handling of complex catalysis, leading to new mathematical results [Sections 2.2, 3]. We then focus on the structure and algorithmic properties of minimal RAFs in [Section 4.1].

## 2 Catalytic reaction systems

### 2.1 Reaction Systems

A *reaction system* is a pair (*X, R*) consisting of a finite nonempty set *X* of *elements* (e.g., molecule types) and a finite set *R* of *reactions*. Here a *reaction r* ∈ *R* refers to an ordered pair (*A, B*) where *A* and *B* are multisets of elements from *X*. We will write *r* : *a*_1_ · · · + → + *a*_*k*_ *b*_1_ + *· · ·*+ *b*_*l*_ to denote the reaction that has reactants {*a*_1_, …, *a*_*k*_ }, and products { *b*_1_, …, *b*_*l*_ } ^1^. We let *ρ*(*r*) denote the set corresponding to *A* (i.e., ignoring multiplicities), and *π*(*r*) denote the set corresponding to *B* (ignoring multiplicities); it is assumed implicitly that *ρ*(*r*), *π*(*r*) ≠ ∅. For a subset *R*′ of *R*, it is convenient to let *π*(*R*′) = ⋃_*r* ∈ *R*′_*π*(*r*) denote the set of the products of the reactions in *R*′.

Next, consider a reaction system (*X, R*) together with a particular subset *F* of *X*. The set *F* can be interpreted as a set of elements that are freely available to the system; accordingly, *F* is referred to as a *food set*. A subset *R*′ is *F-generated* if the reactions in *R*′ can be placed in some linear order, say *r*_1_, *r*_2_, …, *r*_*k*_, such that the following property holds: for *ρ*(*r*_1_) *⊆ F* and for all values of *j* between 2 and *k*, we have *ρ*(*r*_*j*_) ⊆ *F* ∪ *π*({ *r*_1_, …, *r*_*j−*1_ }). In other words, the reactions in *R*′ are F-generated if they can proceed in some order so that the reactant(s) of each reaction are available by the time they are first required. We call such an ordered sequence of *R*′ an *admissible ordering*. Since there are *k*! ways to order *k* reactions, it may not be immediately obvious that the F-generated condition can be verified in polynomial time; however, there is a simple way to do so, as we now describe.

We first recall some further terminology. Given a subset *R*′ of reactions *R*, a subset *W* of *X* is said to be *R*′*-closed* precisely when each reaction *r* ∈ *R*′ that has all its reactant(s) in *W* also has all its the product(s) in *W* (i.e. *r* ∈ *R*′, *ρ*(*r*) ⊆ *W* ⇒ *π*(*r*) ⊆ *W*). The union of two closed sets need not be closed; nevertheless, given any nonempty subset *W*_0_ of *X* there is a unique minimal *R*′-closed set containing *W*_0_, denoted cl_*R*′_ (*W*_0_). This can be computed in polynomial time in the size of the system by constructing a nested increasing sequence of subsets of the elements *W*_0_ ⊂ *W*_1_, … ⊂ *W*_*k*_ where:

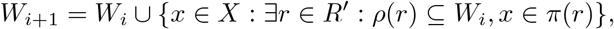

for *i ≥* 0, and by letting *W*_*k*_ denote the terminal set in this sequence (i.e. *k* is the first value of *i* for which *W*_*i*_ = *W*_*i*+1_).

### 2.2 Catalytic reaction systems (allowing complex catalysis)

A *catalytic reaction system* (CRS) is a reaction system with a food set (*X, R, F*) together with a subset *χ* of 2^*X*^ *×R*. Thus *χ* is a collection of pairs (*U, r*) where *U ⊆ X* and *r ∈ R*. For (*U, r*) *∈ χ* we refer to *U* as a *catalyst set* for *r*. This is a generalization of earlier treatments in which *χ* consisted of a subset of *X × R* (i.e. simple catalysis by single elements). Our extension here to this more general catalysis framework allows for complex (i.e. conjunctive) catalysis rules, where catalysts of a reaction may require the presence of two or more elements of *X* (e.g., cofactors of enzymes). However, the treatment of complex catalysis in [41] required the introduction of fictitious new reactions and elements to the original CRS. Here, our more direct approach allows both simple and complex catalysis rules that require no additional reactions or elements to be introduced. It also permits the further option that particular uncatalysed reactions can appear in an autocatalytic system (since the definition of *χ* allows (∅, *r*) ∈ *χ*), thereby addressing a recent concern discussed in Section 2.5.

We denote a CRS by writing 𝒬 = (*X, R, χ, F*), and we let |𝒬| = |*X*| + |*R*| + |*χ*| denote the *size* of 𝒬 and we write

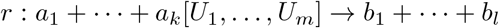

to denote the reaction that has reactants {*a*_1_, …, *a*_*k*_}, products {*b*_1_, …, *b*_*l*_} and catalyst sets {*U*_1_, …, *U*_*m*_}, where (*U*_*i*_, *r*) *∈ χ* for all *i*. When *U*_*i*_ is a singleton set (say {*c*_*i*_}), we will often write *c*_*i*_ in place of {*c*_*i*_} in our example systems.

### 2.3 RAFs

Given a CRS 𝒬 = (*X, R, χ, F*), a subset *R*′ of *R* is said to be *Reflexively-Autocatalytic and Food generated* (more briefly a *RAF*) if *R*′ is nonempty and if for each *r* ∈ *R*′, the reactants of *r* and at least one catalyst set *U* for *r* (as specified by *χ*) is a subset of cl_*R*′_ (*F*).

An equivalent definition for a nonempty set *R*′ to be a RAF for *Q* is that *R*′ is F-generated, and each reaction *r* ∈ *R*′ has a catalyst set *U* that is a subset of *F* ∪ *π*(*R*′). A further equivalent definition is the following:

*R*′ can be ordered *r*_1_, *r*_2_, …, *r*_*k*_ so that for each *i ≥* 1, the reactants of *r*_*i*_ are present in *X*_*i*_, where *X*_1_ = *F* and *X*_*i*_ = *F ∪ π*({*r*_1_, …, *r*_*i−*1_}), and at least one catalyst set *U* of *r*_*i*_ is a subset of *X*_*k*_.

If a CRS 𝒬 has a RAF, then it has a unique maximal RAF (which is the union of all the RAFs for 𝒬), which is denoted maxRAF(𝒬). A RAF *R* for 𝒬 is said to be an *irreducible* RAF (more briefly an *iRAF*) if *R\*{r} is not a RAF for 𝒬 and contains no RAF for 𝒬.

A stronger notion than a RAF is a *CAF* (constructively-autocatalytic and F-generated set) where the third equivalent definition of a RAF (above) is strengthened to ‘and at least catalyst set *U* of *r*_*i*_ is a subset of *X*_*i*_’ (rather than ‘of *X*_*k*_’). In other words, *R*′ is a CAF if it has an admissible ordering in which at least one catalyst set has each of its elements already present in the food set or produced by an earlier reaction in the ordering. Every CAF is also a RAF, but the converse containment does not hold. Although RAFs and CAFs appear to be very similar concepts, they exhibit quite different properties. For example, if a CRS *Q* has a CAF, then this CAF must contain a reaction *r* for which all the reactants of *r* and at least one catalyst of *r* lie in *F*, in which case {*r*} is itself a CAF of size 1. By contrast, a large RAF need not contain any ‘small’ RAF within it. Moreover, theoretical and simulation studies on polymer systems reveal that the level of catalysis required for a CAF to be present is exponentially higher than that required for a RAF [32], and in real biochemical systems that have been studied (e.g., [38, 46]), the maxRAF is generally not a CAF. Thus, in this paper, we focus on the more general notion of a RAF.

### 2.4 Examples

We now describe three examples to illustrate the concepts above. The first example (from [16]) illustrates the concept of a RAF in the simpler setting where catalysis involves only singleton elements, the second example illustrates complex catalysis and the third example is from an experimental system.

Consider the following CRS where *F* = { *f*_1_, *f*_2_, *f*_3_, *f*_4_}, *X* = *F* ∪ { *p*_1_, …, *p*_6_}, and *R*′ = { *r*_1_, …, *r*_6_} indicated by squares in Fig. 1. In this figure, reactants and product pathways are indicated by solids arrows, and catalysis is indicated by dashed arrows.The maxRAF for this system consists of the four reactions (*r*_1_–*r*_4_), and there is one iRAF for this CRS, namely {*r*_1_, *r*_2_}.

**Fig. 1.**
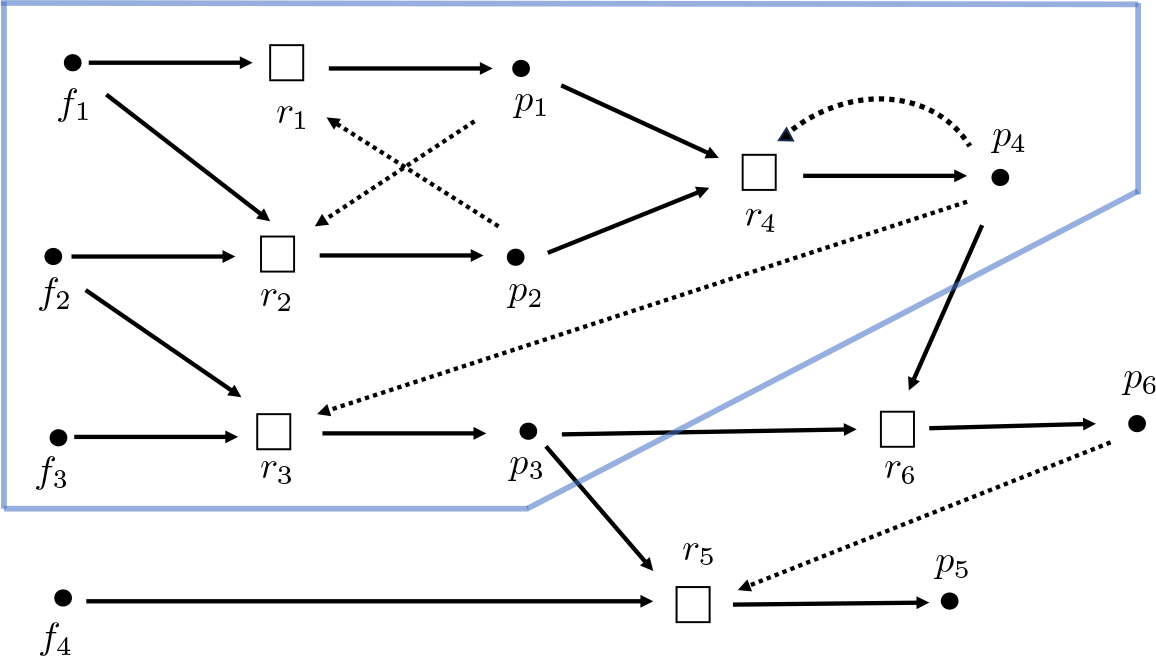
A simple example of a CRS with food set { *f*_1_, *f*_2_, *f*_3_, *f*_4_} six reactions (*r*_1_–r_6_) and with catalysis indicated via dashed arrows (adapted from [16]). The maxRAF consists of the four reactions within the blue border.

Next consider the following CRS where *X* = {*a, b, c, d, e, g*}, *F* = {*a, b*} and *R* = {*r*_1_, *r*_2_, *r*_3_, *r*_4_} and *χ* are as follows:

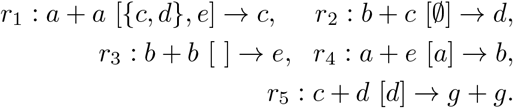

Note the subtle difference between the catalyst set in *r*_2_ and *r*_3_. In *r*_2_ we have *χ* = {∅} so ∅ ∈ *χ* which means *r*_2_ does not require a catalyst (either present in *F* or as a product of another reaction), whereas for *r*_3_, *χ* = ∅ so no catalyst set (*U*) exists for *r*_2_. This system has {*r*_1_, *r*_2_, *r*_5_ } as its maxRAF, and { *r*_1_, *r*_2_} as its unique iRAF. No CAF is present in this CRS.

A third example of a RAF arising in an experimental system (involving simple rather than complex catalysis) is provided in Fig. 2.

**Fig. 2.**
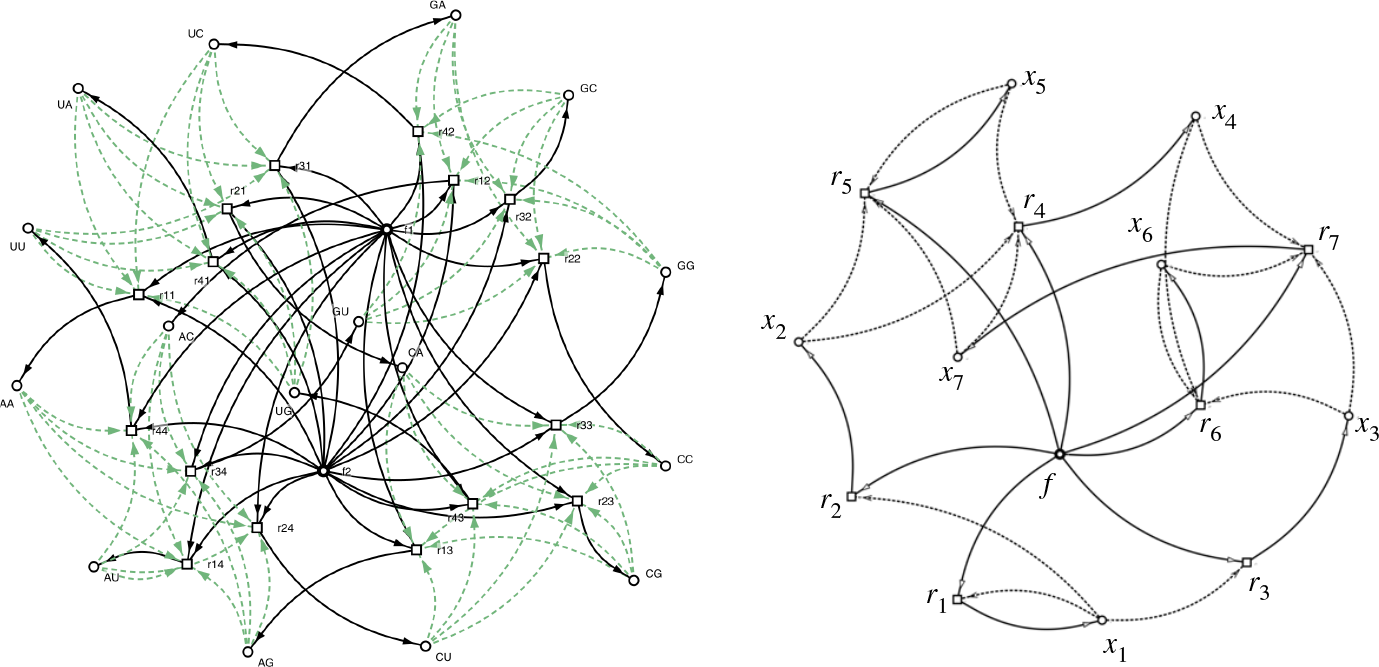
*Left:* An experimental RNA system described in [43] involving 16 reactions, and 18 molecule types (including a food set of size 2, which form the reactants of each reaction). The reactions are denoted by squares, elements (in this case, RNA replicators) are denoted by circles, reaction pathways are indicated by solid arrows, and catalysis is indicated by dotted arrows. This system forms a RAF [16]. *Right:* A subset of seven of the reactions from the full set (with the two food elements combined into a single element *f*), which forms a RAF. This RAF, analysed in [16] contains within it 67 other subsets that form RAFs including four irreducible RAFs (iRAFs). This RAF is not a CAF (nor does it contain one). Figures produced by *CatlyNet* [21].

### 2.5 The interpretation of catalysis and autocatalysis in RAFs

Note that if (∅, *r*) ∈ *χ*, then *r* does not require any element of *X* for its catalysis, and so RAFs under our more general definition can contain uncatalysed reactions in settings where this is appropriate. This is particularly relevant, as a number of papers (e.g., [1, 14, 28, 26]) have pointed to the restrictive nature of RAFs in requiring that all reactions in the RAF must be catalysed. For example, the authors of [1] stated:

> “Here, the assumption that all reactions are catalysed appears very unrealistic. Unfortunately, the algorithms for recognizing RAFs […] do not seem to generalise to arbitrary networks composed of non-catalysed reactions.”

and in [14] the author states:

> “The requirement that every reaction be catalyzed by another molecule seems too strong when we are dealing with simple metabolisms in the earliest forms of life. In modern organisms, almost every reaction of small-molecule metabolites is catalyzed by enzymes. However, prior to the existence of RNA and proteins, there were no enzymes, so we need a theory that deals with the reactions of small molecules without insisting that the reactions be catalyzed.”

In fact, in previous applications of RAF theory to the origins of metabolism [46, 47] a fictional catalyst ‘Spontaneous’ was assigned to reactions known to occur uncatalysed. This catalyst was added to the food set in all simulations. Also, prior to the advent of genetic coding and enzymes, catalysis must have existed in the small-molecule world as well. Small molecules are increasingly being shown to catalyse multiple reactions in the absence of enzymes [3, 33]. However, the use of a fictional catalyst ‘Spontaneous’ is less direct than our approach introduced here (see Section 2.2) where we generalise the notion of catalysis by describing it via a subset *χ* of 2^*X*^ *×R* (thereby allowing pairs of the form (∅, *r*), and so a ‘Spontaneous’ catalyst is no longer required).

RAF theory also treats ‘catalysis’ in a general way and this allows for efficient graph-theoretic algorithms, which apply independently of any detailed kinetic (or even stoichiometric) considerations. This generality also allows for applications in a variety of areas outside chemistry. Essentially, we regard a catalyst as any element that facilitates, speeds-up, or synchronises a reaction without taking part in the reaction itself (as a reactant).

For example, in economic applications, a factory facilitates (i.e. catalyses) the production of items from the incoming raw materials, but is not itself consumed by that process. In cognitive modelling [7], a reaction that combines ideas to form a new idea could be enhanced (i.e. catalysed) by a need, thought, memory, or stimuli. In ecology [9], a catalyst is a species that enables some other interaction in an ecological network. In biochemistry, it can also be helpful to treat catalysis in a quite general way; for instance, the formation of a lipid boundary to encompass a primitive metabolism can be viewed as a sequence of reactions which, once complete, forms a structural element (a complete lipid membrane) that catalyses all the reactions within the newly formed protocell (since the system within it no longer disperses) [17].

Moreover, in biochemistry, food elements can be direct catalysts for multiple reactions, as is the case with universally-essential metal ions. The generality of RAF theory allows for this, in contrast to some other models of network autocatalysis.

It is tempting to treat a catalyst *x* of a reaction *r* as simply an additional reactant and product of *r* (i.e., adding *x* to both sides of the reaction, as is sometimes used in chemical notation), thereby reducing the notion of a RAF to simply an F-generated subset of the resulting (modified) reaction system. In other words, the catalysed reaction *x* + *y*[*c*] → *z* (where *c* is a catalyst) might be viewed as *x* + *y* + *c* →*z* + *c*. However, this misses an important distinction. Namely, in RAF theory, it is assumed that a reaction *r* may proceed (at a slow rate intially) provided that all its reactants are available even if no catalyst is initially available; however *r* will subsequently be part of a RAF set, provided that a catalyst of *r* is (eventually) produced by at least one other reaction in the RAF. Indeed, it is entirely possible for a CRS to contain a RAF yet the modified reaction system (adding catalysts to both sides of a reaction) might have no F-generated subset (indeed, this has been observed in real biochemical networks [16, 38, 46]). Misunderstandings concerning the related notion of ‘autocatalysis’ have been discussed in [18] and [36].

## 3 Mathematical aspects of RAFs

We now recall the concept of the maxRAF operator *φ* (from [42], Section 3). For any subset *R*′ of *R*, let 𝒬|*R*′ be the CRS (*X, R*′, *χ*′, *F*) where *χ*′ is the restriction of *χ* to 2^*X*^ *× R*′, and let *ϕ* : 2^*R*^ *→* 2^*R*^ be the function defined as follows:

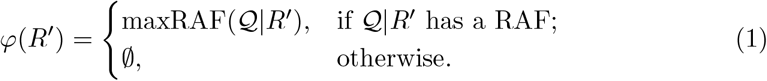

The function *φ*(*R*′) can be determined in a computationally efficient way, as follows:

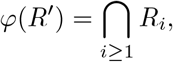

where *R*_1_ = *R*′ and for *i >* 1,

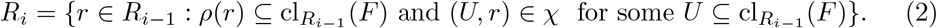

Notice that the sets *R*_*i*_ defined in Eqn. (2) form a decreasing nested sequence, and so *ϕ*(*R*′) is precisely the set *R*_*j*_ for the first value of *j* ≥ 1 for which *R*_*j*+1_ = *R*_*j*_. In particular, a nonempty subset *R*′ of *R* is a RAF if and only if *φ* (*R*′) = *R*′. Moreover, *φ* (*R*) is precisely the maxRAF of 𝒬 (if it exists), or is the empty set otherwise.

**Example:** To illustrate this algorithm, consider the second example described in Section 2.4. For *R*′ = *R* = {*r*_1_, …, *r*_5_} we have *R*_1_ = *R*, and so 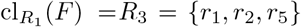, and all but one of the reactions (namely, *r*_3_) satisfy the property that (*U, r*) ∈ *χ* for some 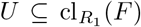. Thus, (by Eqn. (2)), *R*_2_ = {*r*_1_, *r*_2_, *r*_4_, *r*_5_}. Next, *R*_3_ = {*r*_1_, *r*_2_, *r*_5_} since one of the reactants of *r*_4_ ∈ *R*_2_ (namely *e*) is not present in 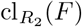. Finally, *R*_4_ = *R*_3_, and so *ϕ*(*R*) = *R*_3_ forms the maxRAF for 𝒬.

A number of interesting algebraic (semigroup) properties of the map *φ* have been explored recently in [29] (see also [30], which considers a more general notion than a RAF, which corresponds to what have been called ‘pseudo-RAFs’ in the RAF literature).

The map *φ* is also an example of the more general notion of an interior operator (on 2^*R*^), and several basic results in RAF theory can be derived by this property alone. To explain this further, an arbitrary function *ψ* : 2^*Y*^ *→*2^*Y*^ is said to be an *interior operator*^2^ if it satisfies the following three properties (nesting, monotonicity, and idempotence) for all subsets *A, A*^*t*^ of *Y* :

(*I*_1_): *ψ* (*A*) ⊆ *A*,

(*I*_2_): *A* ⊆ (*A*)′ ⇒ *ψ* (*A*) ⊆ ψ (*A*)′, and

(*I*_3_): *ψ* (*A*) ⊆ (*A*)) = *ψ* (*A*).

### Lemma 1.

*Given any CRS 𝒬* = (*X, R, χ, F*) *the maxRAF function ϕ* : 2^*R*^ *→*2^*R*^ *is an interior operator*.

Many results of RAF theory (including some we discuss later) depend mainly on the interior operator property of *ϕ*. A natural question is whether *any* interior operator on a finite set can be represented as the maxRAF operator of a set of reactions associated with the elements of the set. It was recently shown that if *Y* is a finite set of size at least 12, then there exist interior operator on *Y* that cannot be represented as a maxRAF operator of a suitably chosen CRS (*X, R*_*Y*_, *χ, F*) where *R*_*Y*_ is a set of reactions bijectively associated with *Y* and where *χ ⊆ X × R*_*Y*_ (i.e. the original CRS setting which does not allow complex catalysis rules) [39]. Thus some generic properties of RAFs in this setting cannot be established by using the interior operator properties alone. However, if one allows complex catalysis rules, we have the following contrasting result (further details and a proof are provided in the Appendix).

### Proposition 1.

*For any finite set Y, any interior operator ψ on* 2^*Y*^ *can be represented as the maxRAF operator ϕ for a suitably chosen CRS 𝒬* = (*X, R*_*Y*_, *χ, F*), *where R*_*Y*_ *is bijectively associated with Y and χ ⊆* 2^*X*^ *× R*_*Y*_ *permits complex catalysis rules. Moreover, R*_*Y*_ *can be chosen so that each of its reactions has the same single reactant f (from the food set) and a single associated product*.

Proposition 1 has an interesting consequence for the structure of RAFs in any CRS. Specifically, it implies that it is possible to simplify the reactions and the food set, while preserving the number, sizes, and containment structure of the RAFs (albeit at the price of possibly increasing substantially the number and complexity of catalysts for the reactions). More precisely, we have the following result (obtained from Proposition 1 by taking *ψ* to be the maxRAF operator of 𝒬).

### Corollary 1.

*For any CRS 𝒬* = (*X, R, χ, F*), *there is a matching CRS 𝒬*′ = (*X*′, *R*′, *χ*′, *F*′), *where (i) each of the reactions in R*′ *has a single reactant f which comprises the food set F, and a single distinct product and (ii) there is a canonical bijective correspondence between R and R*′ *that induces a bijection between the RAFs of 𝒬 and the RAFs of 𝒬′*.

A worked example to illustrate this result is provided in the Appendix. We return to interior operators in Section 4.2.

### 3.1 Strictly autocatalytic RAFs

If a CRS 𝒬 has the property that each reaction is catalysed by at least one element of the food set, then the RAF sets for *𝒬* coincide precisely with the *F* -generated sets. More generally, for any CRS, it is possible for a RAF set to have the property that some (or all) of its reactions are catalysed by elements of the food set, which renders the notion of ‘autocatalytic’ less applicable (though not entirely, since each reaction in the RAF might still also be catalysed by at least one product of the other reactions in the RAF). To formalise this notion further, we introduce a new definition and describe a result that characterises the condition which captures a more focused notion of an ‘autocatalytic’ RAF.

Given a CRS 𝒬 = (*X, R, χ, F*), we say that a subset *R*′ of *R* is *strictly autocatalytic* if each reaction in *R*′ has at least one catalyst that involves one or or more products of reactions of *R*′ and is not a subset of the food set. We say *R*′ is a strictly autocatalytic RAF for 𝒬 if *R*′ is a RAF and is strictly autocatalytic.

**Example:** Consider the following catalytic reaction system:

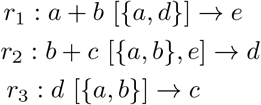

where *X* = {*a, b, c, d, e*} and *F* = {*a, b, c*}. Then, {*r*_2_}, {*r*_1_, *r*_2_}, {*r*_2_, *r*_3_} and {*r*_1_, *r*_2_, *r*_3_} are RAFs, however {*r*_1_, *r*_2_} is the only strictly autocatalytic RAF.

Let 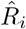, *i ≥* 1 be the nested decreasing sequence of subsets of *R* defined as follows: 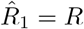, and for *i >* 1, consider the following modification of Eqn. (2):

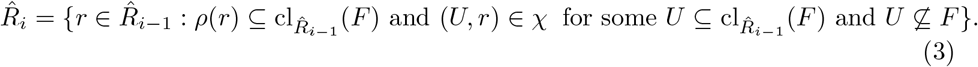

The following result shows that there is a polynomial-time algorithm (in |𝒬|) to determine whether or not 𝒬 has a strictly autocatalytic RAF and, if so, to construct a maximal one. The proof is provided in the Appendix.

#### Proposition 2.

*Le t 𝒬* = (*X, R, χ, F*) *be a CR S. Then 𝒬 has a strictly autocatalytic RAF if and only if* 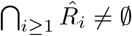, *in which case*, 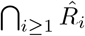 *is the unique maximal strictly autocatalytic RAF for 𝒬*.

#### Application

We investigated the large primitive metabolic data-set used in [46] based on 6039 reactions and a food set of size 68, which resulted in a maxRAF of size 1357. Applying the strictly autocatalytic RAF algorithm determined that no such RAF exists. This, in turn, implies that in any RAF for this system, one or more elements in the food set play a pivotal role in catalysing some reaction(s). This is consistent with the aforementioned essentiality of metals in biocatalysis of central metabolism. At the same time, this result reveals that there are unknown prebiotic routes (or unknown non-enzymatic catalysts) to the production of organic cofactors which must be investigated in the laboratory. For example, there are no routes in this network for the production of organic cofactors as NAD [47] and the investigation of the prebiotic route for its synthesis is subject of active investigation [12, 34].

## 4 Irreducible (minimal) RAFs

Recall that an irreducible RAF (iRAF) is a RAF that contains no RAF as a proper subset. Such RAFs are of particular interest, as they represent the minimal possible autocatalytic networks within a CRS. A smallest sized RAF in any CRS is necessarily an iRAF, however, iRAFs can be of different sizes. Moreover, even without complex catalysis rules, a CRS 𝒬 can have exponentially many iRAFs, and although finding one iRAF is easy, finding a smallest one turns out to be NP-hard [40].

### 4.1 Identifying all iRAFs in a CRS

Let 𝒬 = (*X, R, χ, F*) be a fixed CRS that contains a RAF. Recall that a RAF *R* for 𝒬 is said to be an iRAF (irreducible RAF) if *R\ r* is not a RAF for 𝒬 and contains no RAF for 𝒬. Every RAF of a CRS contains an iRAF, and there is a simple (polynomial-time in |𝒬|) algorithm for finding one or more iRAFs within any given RAF; however, the number of iRAFs for 𝒬 can grow exponentially with |𝒬| (Theorem 1 of [20]). Moreover, determining the size of a *smallest* iRAF is known to be NP-hard [40]. Nevertheless, there is a simple (and polynomial-time) algorithm to determine whether 𝒬 has just one iRAF; we simply ask whether the set

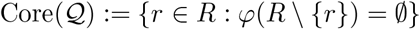

is a RAF. Then it can be shown that this set is a RAF if and only if 𝒬 has a unique iRAF. and in that case, *Core*(𝒬) is the unique iRAF for 𝒬. The result for CRS systems without complex catalysis was established in see [42] Theorem 4.1, however since the proof involved only interior operator properties for *ϕ* it applies to the more general setting here (by Lemma 1).

Moreover, it can easily be seen that there is a simple way to test whether or not an iRAF *R*_1_ is the only iRAF for 𝒬; this holds provided that:

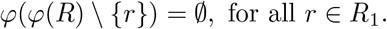

This is computationally less-intensive than computing *Core*(𝒬), since it involves searching over the reactions that are just in *R*_1_ rather than all of *R*.

In a similar way, it can be shown that if 𝒬 has (at least) two iRAFs, *R*_1_ and *R*_2_ then these are the only iRAFs for 𝒬 if and only if for all *r* ∈ *R*_1_ \ *R*_2_, we have *ϕ*(*ϕ*(*R*) { *r* }) = *R*_2_, and for all *r* ∈ *R*_2_ *R*_1_, we have *ϕ*(*ϕ*(*R*) { *r* }) = *R*_1_.

A direct extension of this last result to three or more iRAFs is problematic, since although two iRAFs cannot be nested (i.e. neither can be a subset of the other) in the case of three iRAFs, it is possible for each one to be a subset of the union of the two others. An example is provided by the CRS 𝒬 = (*X, R, χ, F*), where *F* = {*f*}, *R* = {*r*_1_, *r*_2_, *r*_3_} and

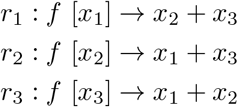

This system has three iRAFs, namely, { *r*_1_, *r*_2_, *r*_1_, *r*_3_}, and { *r*_2_, *r*_3_}, each of which contained in the union of the other two.

However, the following result shows that testing a small number (say, *k*) of iRAFs is feasible in polynomial time in |𝒬| (provided that *k* is fixed). The result is a slight strengthening of Theorem 2 of [40]; its significance is that it allows one to determine whether a given set of *k* iRAFs is the set of *all* iRAFs for 𝒬 in polynomial time in |𝒬|, provided that *k* is fixed (however, the algorithm is exponential in *k*). The proof of the following result is given in the Appendix.

#### Proposition 3.

*A given collection R*_1_, …, *R*_*k*_ *of iRAFs for 𝒬 is the set of* all *iRAFs for 𝒬 if and only if the following condition holds: For all choices* (*r*_1_, …, *r*_*k*_) *where r*_*i*_ ∈*R*_*i*_, *we have ϕ*(*ϕ*(*R*) \{ *r*_1_, …, *r*_*k*_ }) = ∅.

Proposition 3 assumes that the iRAFs are given in advance (e.g., found by heuristic search); however, the approach can be extended to identify the complete set of iRAFs of 𝒬 even if they are not given in advance.

Thus, when the number of iRAF is small (say, less than some value *k*), the following algorithm is polynomial-time in |𝒬| (but is exponential in *k*). The algorithm proceeds as follows and when it terminates, it produces the complete list of iRAFs of 𝒬.

- Given 𝒬 = (*X, R, χ, F*), determine the maxRAF; providing that it exists, compute an iRAF, denoted *R*_1_.
- For *i ≥* 1, construct a sequence of iRAFs starting with *R*_1_, as follows.
  i. For each integer *j* with 1 *≤ j ≤ i* and each choice of *i* reactions {*r*_1_, *r*_2_, …, *r*_*i*_} where *r*_*j*_ ∈ *R*_*j*_ for each *j*, compute *ϕ*(*ϕ*(*R*) \ {*r*_1_, …, *r*_*i*_}). If this set is the empty set for all such choices of {*r*_1_, …, *r*_*i*_}, then *R*_1_, …, *R*_*i*_ is the complete set of iRAFs for 𝒬, so the algorithm terminates.
  ii. Otherwise, if *ϕ*(*ϕ*(*R*) \ {*r*_1_, …, *r*_*i*_}) *≠* ∅ for some choice of {*r*_1_, …, *r*_*i*_}, then compute an iRAF of *ϕ*(*ϕ*(*R*) *r*_1_, …, *r*_*i*_) and set *R*_*i*+1_ equal to this iRAF, and proceed to step (iii).
  iii. Repeat step (i) and, if necessary, step (ii).

### 4.2 Finding an iRAF that contains a given reaction

Given any CRS 𝒬 = (*X, R, χ, F*), a natural question is whether a given reaction in *R* is present in at least one iRAF for 𝒬. We show shortly that this problem is NP-hard. First, we describe how the union of all the iRAFs present in a CRS can be described via a further interior operator. Given a CRS 𝒬 = (*X, R, χ, F*), and a nonempty subset *R*′ of *R*, let 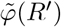 denote the union of the subsets of *R*′ that are iRAFs for 𝒬. The proof of the following lemma is given in the Appendix, as a consequence of a more general result.

#### Lemma 2.

*The map* 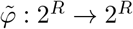 *is an interior operator*.

Although the computation of *ϕ*(*R*′) is polynomial time in |𝒬| for any CRS 𝒬 = (*X, R, χ, F*) and any subset *R*′ of *R*, computing 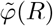 is NP-complete, even for quite simple CRS systems, as we now state more formally (a proof is provided in the Appendix).

#### Proposition 4.

*Given a CRS 𝒬* = (*X, R, χ, F*) *the problem of determining whether or not a given reaction r* ∈ *R is present in at least one iRAF for 𝒬 is NP-complete. Moreover, this holds even for systems were each reaction in R has the same single food-set reactant and where each catalyst consists of single elements (i*.*e. without complex catalysis)*.

Next suppose that a CRS 𝒬 = (*X, R, χ, F*) and that an element *x* ∈ *X \ F* is produced by some reaction in maxRAF(𝒬). A relevant question in certain applications is to find a minimal subset *R*′ of reactions within maxRAF(𝒬) that is both a RAF and produces one or more particular elements of interest (e.g., it plays a key role in a metabolic network, such as ATP or NAD, the universal currencies of energy and redox power in metabolism, respectively). In other words, *R*′ is a RAF that produces *x*, and every proper subset of *R*′ either fails to be a RAF or fails to produce all of the specified elements. It turns out that finding such a minimal set has a polynomial-time solution, as we now show (the proof is provided in the Appendix).

#### Proposition 5.

*Suppose that one or more particular elements x*_1_, …, *x*_*k*_ ∈ *X \ F are produced by some reaction in* maxRAF(𝒬). *Let* 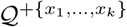 *be the CRS obtained from 𝒬 by replacing each catalyst* (*U, r*) ∈ *χ with* (*U ∪* {*x*_1_, …, *x*_*k*_}, *r*). *Then the collection of minimal subsets of* maxRAF(𝒬) *that are simultaneously RAFs of 𝒬 and produce each of the elements x*_1_, …, *x*_*k*_ *are precisely the iRAFs of* 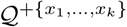.

As an example with the 16-reaction CRS shown in Fig. 2, for each element *x* of *X \ F* the minimal subset that is both a RAF and generates *x* has size either 1 or 2.

### 4.3 Describing a RAF in terms of a composition sequence involving iRAFs

Any RAF contains the union of its iRAFs; however, it may be strictly larger than this union. A simple example is provided by the simple catalytic reaction system:

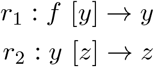

where *F* = { *f* } and *X* = { *f, y, z* }. This system is a RAF but its only iRAF is { *r*_1_ }. In general, a RAF *R*′ is the union of its iRAFs if and only if each reaction in *R*′ is contained in an iRAF.

Nevertheless, any RAF *R*′ in a CRS 𝒬 = (*X, R, χ, F*) can be described by a sequence of iRAFs and associated catalytic reaction systems. This is loosely analogous to the description of finite groups in abstract algebra via a ‘composition series’, in which a group is reduced to the trivial group via quotients that are simple groups.

Here the analogue of a group (respectively, a simple group) is a subset of reactions (respectively, an iRAF), and the analogue of a quotient group is the complement of a iRAF in a set of reactions. However, this analogy is only suggestive; for a finite group the set of associated simple groups is uniquely determined by the group, however, we do not expect the same uniqueness to hold concerning the set of associated iRAFs in our decomposiion.

To describe this in our setting, suppose we have a CRS, 𝒬 = (*X, R, χ, F*) and any nonempty subset *R*′ *⊆ R*. A *composition sequence* for *R*′ is a nested decreasing sequence of subsets of *R*′, *R*_1_, …, with *R*_1_ = *R*′ and with 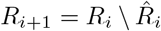 where 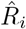 is an iRAF of 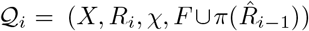 for each *i ≥* 1 for which 𝒬_*i*_ has a (nonempty) RAF. Since the sets *R*_*i*_ form a nested decreasing sequence of sets, we refer to the final distinct set (i. e. ⋂_*i ≥*1_ *R*_*i*_) as the *terminal set* of the sequence.

It is clear that every subset *R*′ of *R* has a composition sequence, and that this can be constructed in polynomial time in |𝒬|. Moreover, by definition, the iRAF sets 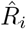 are disjoint. In the simple 2-reaction example above, the unique composition sequence for *R*′ = { *r*_1_, *r*_2_}is: { *r*_1_, *r*_2_}, { *r*_2_},∅. The proof of the following result is provided in the Appendix.

#### Proposition 6.

*Let Q* = (*X, R, χ, F*) *be a CRS, let R*′ *be a nonempty subset of R, and let R*_1_, *R*_2_, … *be a composition sequence for R*′. *The following are equivalent:*

i. *R*′ *is a RAF for 𝒬;*
ii. *R*_1_, *R*_2_, … *has terminal set* ∅.
iii. *The associated iRAF sets* (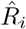, *i ≥* 1) *partition R*′.

Note that the iRAF sets 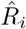 in (iii) are not, in general, iRAFs of the original CRS 𝒬, and since the sets 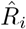 are (by definition) disjoint, the partitioning condition in (iii) is equivalent to the condition that the union of the sets 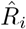 is equal to *R*′.

## 5 Concluding comments

Autocatalytic networks have provided a formal tool to investigate processes in early biochemistry and certain other settings that involve the formation and evolution of complex structures. The discrete nature of the model provides a way to develop and implement efficient algorithms that can be applied to large data sets, and to elucidate their structural properties, such as their building blocks in terms of iRAFs or the ordering of reactions.

In this paper, we have developed further techniques that open the door to more detailed investigations into the structural properties of RAFs and iRAFs, their extensions to more general catalytic scenarios, and the investigation of the constraints on the order of reactions and the appearance of particular elements of interest. Several of the techniques described here have been implemented in the open-source software package *CatlyNet* [21], which we plan to apply to investigate further aspects of early metabolism and related evolutionary questions.

## 6 Acknowledgements

We thank the Royal Society Te Apārangi (New Zealand) for funding under the Catalyst Leader programme (agreement No. ILF-UOC1901). We also thank the two anonymous reviewers for numerous helpful comments and suggestions on an earlier version of this manuscript.

## 7 Appendix Mathematical details

*Proof of Lemma 1*: For a subset *R*′ of *R*, let

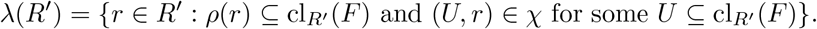

It is clear that the function *λ* satisfies properties (*I*_1_) and (*I*_2_) (but not necessarily (*I*_3_)), and since *ϕ*(*R*′) = ⋂_*i≥*1_ *H*_*i*_(*R*′) where *H*_1_(*R*′) = *R*′ and *H*_*i*+1_(*R*′) = *λ*(*H*_*i*_(*R*′)) for all *i ≥* 1, Property (*I*_3_) holds by Proposition 1 of [39].

*Proof of Proposition 1*: Let *ψ* be an arbitrary interior operator on 2^*Y*^. We construct a CRS 𝒬_*ψ*_ = (*X, R*_*Y*_, *χ, F*) that permits complex catalysis rules, as follows. Let *F* = {*f*}, let *X* = {*x*_*y*_ : *y ∈ Y* } ∪ {*f*}, and let *R*_*Y*_ = {*r*_*y*_ : *y ∈ Y* } where *r*_*y*_ is the reaction *f → x*_*y*_. It remains to describe, for each reaction *r*_*y*_ ∈ *R*_*Y*_, its associated catalyst sets (i.e. the subsets *U* of *X* for which (*U, r*_*y*_) ∈ *χ*). Let

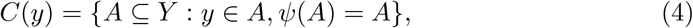

the fixed sets of *ψ* that contain *y*. Then we let (*U, r*_*y*_) be an element of *χ* if and only if *U* = {*x*_*a*_ : *a ∈ A*} for some set *A ∈ C*(*y*).

Let *b* : *Y → R*_*Y*_ denote the bijection *y → r*_*y*_.

### Claim 1

A nonempty set *A* of *Y* is a fixed set of *ψ* if and only if *b*(*A*) = {*r*_*y*_ : *y ∈ A*} is a RAF for 𝒬_*ψ*_.

To establish this claim, suppose first that *A* is a fixed set of *ψ* (i.e. *ψ*(*A*) = *A*). Since any nonempty subset of *R*_*Y*_ is *F* -generated (because each reaction in *R*_*Y*_ has only the single reactant *f* which lies in the food set), it suffices to show that each reaction *r*_*y*_ in *b*(*A*) has an associated catalyst set *U* with the property that each of its elements is produced by at least one reaction in *b*(*A*). Suppose that *r*_*y*_ ∈ *b*(*A*). Then *y ∈ A*, and, by assumption, *A* is a fixed set of *ψ*, so *A ∈ C*(*y*); moreover, the elements {*x*_*a*_ : *a ∈ A*} are products of reactions in *b*(*A*). Thus, taking *U* = {*x*_*a*_ : *a ∈ A*} we have (*U, r*_*y*_) ∈ *χ*; in other words, *r*_*y*_ is catalysed by products of reactions in *b*(*A*). Since this holds for all *r*_*y*_ ∈ *b*(*A*), it follows that *b*(*A*) is a RAF.

Conversely, suppose that *b*(*W*) is a RAF for 𝒬_*ψ*_ for some subset *W* of *Y*. Then for each *w ∈ W*, the reaction *r*_*w*_ : *f → x*_*w*_ has a catalyst set of the form {*x*_*a*_ : *a ∈ A*_*w*_} where *A*_*w*_ is a nonempty subset of *W* satisfying *w* ∈ *A*_*w*_ and *ψ*(*A*_*w*_) = *A*_*w*_. It remains to show that *ψ*(*W*) = *W*. Since *ψ*(*W*) ⊆*W* it suffices to determine the converse containment. We have:

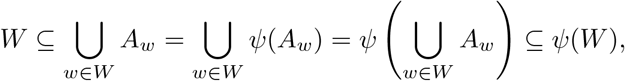

as required to establish Claim 1.

Next observe that the RAFs for 𝒬_*ψ*_ are simply the nonempty fixed sets of *ϕ* (the maxRAF operator on *R*_*Y*_ for the CRS 𝒬_*ψ*_), and so *ψ* and *b*^*−*1^ *◦ ϕ ◦ b* are two interior operators on 2^*Y*^, and these two functions the same fixed sets by Claim 1 (noting also that *ϕ* and *ψ* also trivially fix ∅). However, any two interior operators on 2^*Y*^ that have the same collection of fixed sets are identical functions. This is because any interior map *ν* on a set 2^*Y*^ is completely determined by its fixed sets since for any set *Y* ′⊆ *Y* we have:

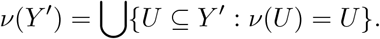

Thus *ψ* = *b*^*−*1^ *◦ ϕ ◦ b*, completing the proof. 0

**Remark:** It can be shown that Claim 1 also holds if we restrict the catalyst sets *U* = {*x*_*a*_ : ∈ *a A*} to just those that correspond to the *minimal* sets *A ∈ C*(*y*) (where minimal refers to set inclusion).

*An example to illustrate Corollary 1*: Consider the following CRS (based on Kauffman’s binary polymer model with food set *F* = {0, 1, 00, 01, 10, 11} from Fig. 1 of [41]):

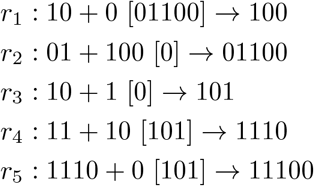

This system is itself a RAF and it contains six other RAFs as subsets, namely {*r*_1_, *r*_2_}, {*r*_3_}, {*r*_1_, *r*_2_, *r*_3_}, {*r*_3_, *r*_4_}, {*r*_3_, *r*_4_, *r*_5_}, {*r*_1_, *r*_2_, *r*_3_, *r*_4_}. Following the procedure described in the proof, we associate with *r*_*i*_ the reaction *rit* : *f* [*U*_*i*_] *→ x*_*i*_ where the set of (complex) catalysts is *U*_*i*_ = {{*x*_*j*_ : *r*_*j*_ ∈ *A*} : *A* is a RAF containing *r*_*i*_}. For example, for *r*_1_ we have:

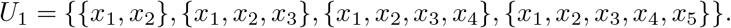

We can further simply *U*_1_ in line with remark following the proof of Proposition 1 above. For *U*_1_ we only need to keep the single set {*x*_1_, *x*_2_} since all other sets in *U*_1_ contain this set. Applying this simplification to the other sets *U*_*i*_ leads to the resulting reduced system:

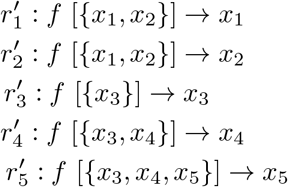

The RAFs of this system (with *F* = {*f*}) then correspond bijectively to the RAFs of the original system (under the bijection 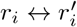 for each *i*).

*Proof of Lemma 2*: We establish the following more general result from which Lemma 2 follows by taking *ψ* to be the maxRAF operator (*ϕ*) for a CRS.

Given any function *ψ* : 2^*Y*^ *→* 2^*Y*^ define an associated derived function 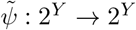 as follows: For *X* ⊆*Y*, let *F*_*ψ*_(*X*) = {*U* ⊆*X* : *ψ*(*U*) = *U* } (the ‘fixed sets’ of *ψ*) and set:

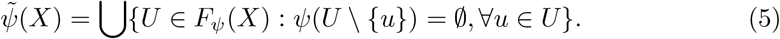

In words, 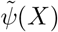 is the set of elements in *X* that lie in at least minimal fixed set of *ψ*.

### Lemma 3.

*If is an interior operator on* 2^*Y*^ *then* 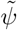 *is also an interior operator on* 2^*Y*^, *and* 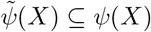 *for all X ∈* 2^*Y*^.

*Proof:* Condition (*I*_1_) holds for 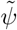 since *ψ*(*X*) *⊆ X* for all *X ∈* 2^*Y*^ and so *U ∈ F*_*ψ*_(*X*) implies that *U ⊆ X*, and by definition 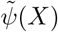 is a union of sets in *F*_*ψ*_(*X*). Condition (*I*_2_) holds for 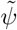 since if *X ⊆ X*′ and *U ∈ F*_*ψ*_(*X*) then *U ∈ F*_*ψ*_(*X*′). Thus it remains to establish (*I*_3_). We have 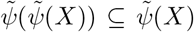 by condition (*I*_1_), and so it suffices to establish the reverse containment. Suppose that *U ∈ F*_*ψ*_(*X*). Then since *U ⊆ X* and *ψ*(*U*) = *U*, it follows that *U* = *ψ*(*U*) *⊆ ψ*(*X*). Thus,

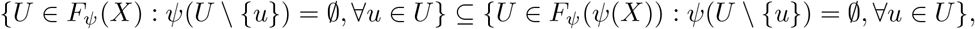

and so the union of the sets on the left (i.e.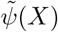) is a subset of the union of the sets on the right (i.e.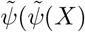)). Thus, by Eqn. (5), (*I*_3_) holds for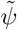, and so 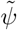 is an interior operator on 2^*Y*^.

The containment 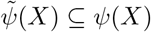 for all *X* ∈ 2^*Y*^ follows by observing from Eqn. (5) that 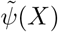 is a union of a sub-collection of the set of subsets of *X* that are fixed by *ψ*, and the union of all such *ψ*–fixed subsets of *X* is *ψ*(*X*). 0

*Proof of Proposition 2*: Suppose that 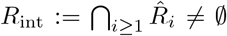. Then *R*_int_ is a RAF, and every reaction of *R*_int_ has a catalyst *U*_*i*_ that is not a subset of *F*, and so *R*_int_ is a strictly autocatalytic RAF. On the other hand, if *R*′ is a strictly autocatalytic RAF, then since *R*′ *⊆ R* we have (by induction on *i ≥* 1) 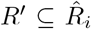 for each *i*. Thus, since 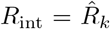 for some value of *k* we have *R*′ *⊆ R*_int_ and so *R*_int_ is the unique maximal strictly autocatalytic RAF for 𝒬.

*Proof of Proposition 3*: The argument just relies on the interior operator properties of *ϕ* established in Lemma 1. If *ϕ*(*ϕ*(*R*) *\ {r*_1_, …, *r*_*k*_}) *∅* then *ϕ*(*R*) *\ {r*_1_, …, *r*_*k*_} contains an iRAF, and this iRAF cannot be one of *R*_1_, …, *R*_*k*_, since for each *i, r*_*i*_ was chosen from *R*_*i*_ yet *r*_*i*_ ∉ *ϕ*(*ϕ*(*R*) *\ {r*_1_, …, *r*_*k*_}) by definition. Conversely, if there is an iRAF *R*′ of *R* that is different from *R*_1_, …, *R*_*k*_, then selecting *r*_*i*_ in *R*_*i*_ *\ R*′ for each *i* (this is possible since *R*_*i*_ is not a subset of *R*′), *R*′ is an iRAF of *ϕ*(*R*) \ {*r*_1_, …, *r*_*k*_} and so *R*′ = *ϕ*(*R*′) *⊆ ϕ*(*ϕ*(*R*) \ {*r*_1_, …, *r*_*k*_}), thus this set is nonempty.

*Proof of Proposition 4*: First note that the problem described lies in the complexity class NP, since it can readily be verified (in polynomial time in |𝒬|) whether or not a given subset of reactions is an iRAF for 𝒬 that contains a given reaction. We reduce an instance of the problem described to the following graph-theoretic problem: Given a finite directed graph *G* = (*V, A*), and a vertex *v* ∈*V*, determine whether or not there exist a chordless cycle in *G* that contains *v*. This problem was shown to be NP-complete in [31] (Theorem 1).

Given an arbitrary finite directed graph *G* = (*V, A*), consider the following CRS 𝒬_*G*_ = (*X, R, χ, F*) where *F* = {*f}, X* = *V ∪ F, R* = {*r*_*v*_ : *v ∈ V* } where *r*_*v*_ is the reaction *r*_*v*_ : *f* [*u*_1_, …, *u*_*n*_] *→ v* for each ordered pair (*u*_*i*_, *v*) ∈*A*. Notice that 𝒬_*G*_ is a CRS for which there is a single reactant for each reaction (namely *f*) which comprises the food set. Moreover, each reaction has singleton catalysts (i.e. complex catalysis is not involved).

Now any CRS 𝒬 = (*X, R, χ, F*) that has the property that (i) each of its reactions has all its reactants in the food set, and (ii) each catalyst of each reaction is a singleton element in *X \ F*, then the iRAFs for 𝒬 correspond precisely to the subsets of *R* that form a chordless cycle of the graph *G*_𝒬_ := (*R, A*) where (*r, r*′) ∈ *A* if a product of *r* is a catalyst of *R*′, (Theorem 2.1(ii) of [41]). It follows that the iRAFs for _*G*_ are precisely the chordless cycles of 𝒬_*G*_, which establishes the claimed reduction.

*Proof of Proposition 5*: Suppose that *R*′ is a minimal subset of maxRAF(𝒬) that is a RAF and produces all of the elements *x*_1_, …, *x*_*k*_. Then *R*′ is a RAF of 𝒬^+^ := 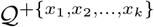. Moreover, if *R*′ is a strict subset of *R*′ that is a both a RAF and that produces *x*_1_, …, *x*_*k*_, then *R*′ is also a RAF of 𝒬^+^ and so *R*′ is not a minimal set with the joint properties of being a RAF and producing *x*_1_, …, *x*_*k*_. Thus *R*′ is an iRAF of 𝒬^+^. Conversely, suppose that *R*′ is an iRAF of 𝒬^+^. Then *R*′ is a RAF of 𝒬 and *R*′ must produce all of the elements *x*_1_, …, *x*_*k*_ (since all reactions in 𝒬^+^ require each of the elements *x*_1_, …, *x*_*k*_ to be produced because none of these elements lie in the food set *F* (by assumption)). Moreover, *R*′ is minimal subset of *R* that is a RAF for and produces *x*_1_, …, *x*_*k*_; for otherwise *R*′ contains a strict subset with these properties and so would be a RAF of 𝒬^+^ contradicting the assumption that *R*′ is an iRAF of 𝒬^+^.

*Proof of Proposition 6*: We begin with a lemma.

### Lemma 4.

*Suppose that R*′ *and R*′ *⊊ R*′ *are both RAFs for* = (*X, R, χ, F*). *Then R*′ *\R*′ *is a RAF for* (*X, R, χ, F∪ π*(*R*′)).

To see why this holds, simply observe that if *o*′ is an admissible ordering for *R*′ and *o*^*−*^ is the induced ordering of *R*′ *\R*′′ then *o*^*−*^ is an admissible ordering of *R*′ *\R*′ for the CRS with expanded food set (*X, R, χ, F∪ π*(*R*′)).

Returning to the proof of Proposition 6, observe that (ii) and (iii) equivalent (by the comments following the last lemma) so it suffices to show that (iii) implies (i), and, conversely, that (i) implies (iii). Suppose first that (iii) holds. Since *R*′ equals the (disjoint) union of the sets 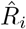 we can select an admissible ordering of 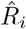 for each *i*, and extend this to an ordering of the reactions of *R*′ in which all reactions from 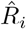 come before those of 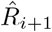 for each *i*. This ordering is an admissible ordering of *R*′, and each reaction in *R*′ is either catalysed by an element of the food set or by the product of another reaction from *R*′. Thus *R*′ is a RAF for 𝒬.

The proof of the converse direction ((i) implies (iii)) follows from Lemma 4, by induction on the length of the sequence *R*_1_, *R*_2_, … and the fact that every RAF contains an iRAF.

## Footnotes

1 In applications, a bidirectional reaction is generally regarded as a pair of reactions (forward and backward) with the same catalysis assignment.

2 The term ‘interior operator’ comes from topology, since the function that assigns to any subspace *S* of a topological space the topological interior of *S* satisfies the three properties described.

## Notes

### Competing Interest Statement

The authors have declared no competing interest.

### Summary of Updates

The original paper has been revised following peer review, with two earlier sections deleted, and other sections restructured and expanded.

